# Highly Efficient Multiplexed Genome Engineering and Clone Selection to Enable Next Generation Induced Pluripotent Stem Cell (iPSC)-based Cell Therapies

**DOI:** 10.64898/2025.12.23.696205

**Authors:** Kaivalya Molugu, Ruba Halaoui, Mohammed Zeaiter, Anne M. Bara, Matthew Bakalar, Bruna Castro de Oliveira, Connor McGinnis, Jie Wang, Marie-Joe Kimaz, Lauren Herchenroder, Arun Bhattathiri, Alexander Allen, Minggang Fang, Davood Norouzi, Didac Santesmasses, Dane Z. Hazelbaker, Daniel J. O’Connell, Dennis Asante, Jason T. Gatlin, Xiarong Shi, Devin Harrison, Sandeep Kumar, Ravindra Amunugama, Jonathan D. Finn, Chong Luo

## Abstract

Human induced pluripotent stem cells (iPSCs) have revolutionized regenerative medicine and cellular therapies. To improve the functionality and safety of iPSC-based therapies, genome engineering has been employed to disrupt gene expression and introduce therapeutic transgenes. However, current genome editing methods face significant challenges, including labor-intensive procedures, low efficiency, and safety concerns. Here, we report a novel and efficient approach for multiplex genome engineering in iPSCs using Integrase-mediated Programmable Genomic Integration (I-PGI) technology. I-PGI combines CRISPR-mediated genome editing and site-specific integrases, enabling precise gene insertion without DNA double-strand breaks. By optimizing I-PGI components and protocols, we achieved multiple gene knockouts and knock-ins of up to five transgenes in a single process. Additionally, we developed a streamlined workflow for enrichment, deposition, and screening of single-cell clones with the desired genome edits. This approach significantly accelerates and improves the precision of multiplex iPSC engineering, paving the way for next-generation iPSC-based therapies.

## Introduction

Induced pluripotent stem cells (iPSCs) have emerged as a transformative platform in regenerative medicine and cellular therapies^1,2^. These cells are derived from somatic cells that have been reprogrammed to a pluripotent state, enabling them to differentiate into virtually any cell type of the human body^3^. iPSCs possess the unique ability to self-renew, making them a renewable and potentially cost-effective source of cells. By using the patient’s own cells, iPSCs enable personalized cell therapies that are genetically matched to the patient^4^. Alternatively, iPSCs derived from healthy donors can be used as off-the-shelf cell products. iPSC-based therapies hold promise for treating a wide range of diseases, from neurodegenerative disorders, cardiovascular diseases, genetic disorders and cancer^5–7^.

Genetic engineering to disrupt target gene expression (knockouts, KOs) and to introduce expression of therapeutic transgenes (knock-ins, KIs) have been actively explored by researchers to enhance the functionality and safety of iPSC-based therapies. Examples of engineering strategies include stealth edits to evade immune recognition, and suicide switches to enable the eradication of any undifferentiated iPSCs and the removal of cell products when adverse events occur in patients^8,9^. We have detailed 12 engineering strategies used in iPSC-derived immunotherapies in our recent review^10^.

Advanced genome editing technologies are needed to support sophisticated engineering of iPSCs. Over the past decade, CRISPR/Cas9-based technologies have revolutionized basic and applied research in biology. However, current gene integration approaches generally require DNA double-strand breaks (DSBs), which risk chromosomal translocations, and rely on repair pathways such as homology-directed repair (HDR) that are inactive in quiescent or terminally differentiated cells. Achieving programmable and multiplexed genome integration of multi-kilobase DNA cargo in an HDR-independent manner therefore remains a significant challenge. Building on these advances, CRISPR/Cas systems have increasingly been coupled with established families of editing enzymes, such as transposases, integrases, recombinases, leading to the development of new technologies capable of long sequence integration^11^.

Here, we report a highly efficient, multiplexed genome editing process in iPSCs using the Integrase-mediated Programmable Genomic Integration (I-PGI) technology. I-PGI is derived from the Programmable Addition of Site-Specific Targeting Elements (PASTE) technology and is mechanistically related to PASTE and PASSIGE^12,13^. These platforms all use a Cas9 nickase (nCas9) fused to a writing enzyme (e.g., reverse transcriptase) together with attachment site–containing guide RNAs (atgRNAs) to install a large serine integrase (LSI) landing site, herein referred to as a “beacon,” at a specific location in the genome. Following beacon placement (BP), expression of the corresponding LSI, such as Bxb1 integrase, along with a template DNA containing the cognate recognition site, leads to the targeted integration of the template DNA at the beacon^13^. LSIs are agnostic to the size of the template DNA, and mediate irreversible, unidirectional recombination independent of host cell DNA repair mechanisms, therefore enabling activity in both dividing and quiescent cells^14^. I-PGI thus enables precise and efficient multiplex genome editing in iPSCs without DSBs.

We aimed to develop an allogeneic iPSC-derived natural killer (iNK) cell product using I-PGI to reset the humoral immune system in patients with autoimmune diseases. To this end, we introduced a number of KOs and KIs into iPSCs, including disruption of class I and II Human Leukocyte Antigens (HLAs) (β2-microglobin, B2M and CIITA KOs) to evade allogeneic immune recognition; elimination of CD52 expression to enable antibody-mediated lymphodepletion; and expression of the following transgenes: 1) a CD19/BCMA dual-targeting chimeric antigen receptor (CAR) for antigen-specific cytotoxicity; 2) IL-15/IL-15Rα to enhance NK cell metabolism and persistence; 3) an inducible Caspase 9 (iCasp9) safety switch for rapid elimination of the cells when needed; and 4) a synthetic HLA-E and B2M fusion protein to evade NK-mediated immune rejection. Gene knockouts of B2M, CIITA and CD52 were achieved by placing beacons with orthogonal dinucleotide sequences to disrupt the endogenous protein expression. DNA cargos encoding the transgenes were subsequently integrated into the B2M beacon alone, or into both B2M and CIITA beacons. The orthogonal beacons at CIITA and CD52 provide flexibility for future engineering, as they can be used to insert additional cargo into the same iPSC clone.

Following genome engineering using I-PGI in iPSCs, we developed a streamlined workflow to enrich gene-edited cells during single cell deposition, and to identify colonies with the intended edits using high-throughput next-generation amplicon sequencing. The selected iPSC clones can then be differentiated into cell types of interest and subjected to downstream functional assessment. Together, we here describe a novel approach to engineer and select multiplex genome-edited iPSCs with the advantages of speed and precision, accelerating the development of future complex edited cell therapies.

## Materials & Methods

### iPSC culture & passaging

Human iPSCs were obtained from ATCC (ACS-1025) and Fujifilm Cellular Dynamics, Inc. iPSCs were maintained in E8 medium (Cat A1517001, ThermoFisher Scientific) on vitronectin (A31804, ThermoFisher Scientific)-coated tissue culture polystyrene plates (Corning). Cells were passaged every 4–5 days at a ratio of 1:10 using ReLeSR solution (STEMCELL Technologies). All cells were maintained at 37 °C in 5% CO_2_ and tested monthly for mycoplasma contamination.

### Electroporation for Beacon Placement and Programmable Genomic Integration

iPSCs were singularized using Accutase (Cat 07922, STEMCELL Technologies) and counted using Via2-Cassette (Cat 941-0024, Chemometec) in the NC-200 (Chemometec) instrument. iPSCs were then seeded at 35,000 cells/cm^2^ on 6-well plates (Corning) in 2 mL of E8 media and 10 μM ROCK inhibitor (cat S1049, Selleckchem), 48 hours before electroporation. iPSCs were electroporated using the Neon Transfection System 10 μL kit (Cat MPK1096, ThermoFisher Scientific) as per the manufacturer’s instructions. Briefly, iPSCs were harvested using Accutase (STEMCELL Technologies) and counted. 1 × 10^5^ cells per electroporation were then centrifuged at 300 g for 5 min. Media was aspirated and cells were resuspended using 10 μL of R buffer (ThermoFisher Scientific) with 1) 50 pmol each of dual atgRNA for the desired site and 1 pmol of nCas9-RT mRNA for beacon placement or 2) 1 μg of desired DNA donor cargo and 1.5 pmol of Bxb1 integrase mRNA. iPSCs were then electroporated using 1200 V,30 ms,1 pulse for Beacon Placement and 1500 V, 20 ms,1 pulse for Programmable Genomic Integration. After electroporation, cells were plated into 48-well plates in 100 μL of E8 media + 1X CloneR2 (Cat 100-0691, STEMCELL Technologies). Media was changed 24 h post-transfection and replaced with E8 medium daily. After incubating at 37 °C, cells were harvested for ddPCR/NGS analysis on Day 3 and for flow cytometry analysis on Day 7.

### Genomic DNA extraction

Genomic DNA (gDNA) was extracted 3 days after electroporation following treatment with Accutase, centrifugation, and resuspending cells in 50 μL QuickExtract (Cat QE0905T, LGC Biosearch Technologies) following manufacturer’s instructions. Briefly, cells in QuickExtract were incubated at 65 °C for 6 min followed by 98 °C for 2 min. gDNA was then purified from cell lysates by AMPure XP magnetic bead (Cat A63882, Beckman Coulter) according to manufacturer’s instructions.

### Target amplification and next-generation amplicon sequencing (NGS)

Target regions were amplified with Q5 Hot Start High-Fidelity Master Mix (M0494X, NEB) for 25 cycles using annealing temperatures for the gene-specific part of the primers calculated by NEB’s online tool (https://tmcalculator.neb.com/). Amplified targets were either imaged with gel electrophoresis in a 2% agarose gel or used as a template for the Illumina barcoding PCR 2.

For rhAmpSeq analysis of off-target edits, off-target sites for each atgRNA pair were predicted by CALITAS^17^ and PCR panels were ordered from IDT and performed according to manufacturer’s instructions.

### Digital droplet PCR (ddPCR)

Custom primers and probes were designed to measure beacon placement and programmable genomic integration in iPSCs. Results were normalized to custom reference assays targeting unedited regions of the same genes. Probes were dual labeled with 3′-3IABkFQ and either 5′-carboxyfluorescein (FAM) for edit targets or 5′-hexachlorofluorescein phosphoramidite (HEX) for reference. Assays were validated using gBlocks representing edit outcomes to test for both specificity and linearity. All primers, probes, and gBlocks were synthesized by IDT (Coralville, IA, USA). The reaction mix for all reactions, was composed of 12 µL of 2x□ddPCR Supermix for probes (No dUTP) (Cat 1863025 Bio-Rad), 1.2 µL of each primer and probe mix. 1 µL genomic DNA and water to a final volume of 25 µL. Droplets were generated on the AutoDG Instrument for automated droplet generation (Cat 186410, Bio-Rad, Hercules, CA, USA). PCR amplification was performed with the following cycling parameters: initial denaturation at 95 °C for 10 min, followed by 40 cycles of denaturation at 94 °C for 30 sec and combined annealing/extension step at 58 °C for 1 min, and a final step at 98 °C for 10 min, ending at 4 °C, all with controlled temperature ramps of 1°C/sec. Data acquisition and analysis were performed on the QX200 Droplet Reader using the QX Manager software (Cat 1864003, Bio-Rad).

### Flow Cytometry

Cells were collected in a 96 well plate, washed twice with Cell Staining Buffer (BioLegend #420201). Antibodies and Live/Dead Stain were added at 1:50 and 1:100 dilutions, respectively in the Cell Staining Buffer and incubated for 20-30 min at 4 °C. This was followed by another 2 washes in Cell Staining Buffer and a final resuspension in 100 μL. The samples were then run on the CytoFlex LX flow cytometer (Beckman Coulter).

| Target | Catalog Number | Company | Dilution |
| --- | --- | --- | --- |
| rhCD19 | CD9-H52H2 | ACRO | 1:50 |
| rhBCMA | BCA-HA2H9 | ACRO | 1:50 |
| IL-15Rα | 747699 | BD Biosciences | 1:50 |
| HLA-E | 53-9953-42 | Invitrogen | 1:50 |
| HLA-E | 342610 | BioLegend | 1:50 |
| HLA-E | 342604 | BioLegend | 1:50 |
| HLA-E | 342608 | BioLegend | 1:50 |
| LIVE/DEAD | L34980 | Life Technologies | 1:100 |

### Single cell deposition

Cells were passaged, washed and stained using a 1:50 antibody dilution, incubating at room temperature for 30 min. Cells were then washed twice and counted. Final resuspension was adjusted to 10,000-20,000 cells/ml. 600 μL of the cell suspension was loaded into the Namocell Pala cartridge and cell analysis was run on the machine. A gate was drawn over the live cell population using the FSC/TSC plot, and the fluorophore positive population in the fluorophore/TSC plot. The selected two gates will determine which cells will be deposited into the 96-well plates. Single cell sorted deposition was launched into 96-wel lplates previously loaded with 100 μL of media/well with 10% CloneR2. Once deposition of the plate is completed, the plate is moved to the 37 °C incubator.

### Single cell imaging

Following cell deposition, the plates were centrifuged at 400 ×g for 5_min, thereby ensuring that all cells settled at the bottom of the wells. Immediately after centrifugation, wells of the 96-well plates were imaged using a Cellavista imager (Synentec, Germany)^15^. Plates were incubated at 37 °C in a humidified atmosphere with 5% CO_2_ for outgrowth. Confluence per well in 96-well plates was measured using the same imager. The results were analyzed using YT-software (Synentec, Germany).

### Karyotyping

Cells cultured for at least five passages were grown to 60– 80% confluence and shipped for karyotype analysis to WiCell Research Institute, Madison, WI. G-banded karyotyping was performed using standard cytogenetic protocols. Metaphase preparations were digitally captured with Applied Spectral Imaging software and hardware. For each cell line, 20 GTL-banded metaphases were counted, of which a minimum of 5 were analyzed and karyotyped. Results were reported in accordance with guidelines established by the International System for Cytogenetic Nomenclature 2016.

### Statistical analysis

Graphs and data analysis were performed with GraphPad Prism 10. All results are reported as mean and standard deviation of biological duplicates or triplicates with standard deviation for all the experiments.

## Results

### Beacon placement in iPSCs

Our I-PGI technology uses nCas9, reverse transcriptase, and a pair of atgRNAs with partially overlapping beacon sequences, to insert a full beacon for cargo integration by the large serine integrase Bxb1. For guide RNA spacer sequence selection, we utilize CRISPOR^16^, which identifies guide RNAs in a given sequence and ranks them based on various scores that assess potential off-target effects in the genome and predict on-target activity. Off-target effects are ranked using CFD and MIT specificity scores, while a more detailed off-target profile is obtained through CALITAS^17^.

After excluding atgRNAs with significant off-target potential, we predict on-target editing efficiencies using our proprietary data-driven model. Previous studies have demonstrated that the editing efficiencies of similar single prime editors can be accurately predicted based on the guide RNA sequence composition, structural parameters, and local chromatin accessibility^18,19^. Similarly, we developed a machine learning model to identify the factors that influence beacon insertion frequency and fidelity. The key factors include the GC content and melting temperatures of the spacer and primer binding sites (PBSs), nicking distance, open chromatin signals (e.g., ATAC-seq), heterochromatin markers (e.g., H3K9me3), and atgRNA structural features such as PBS length, scaffold type, partial beacon overlap size, and guide RNA modification patterns. Based on these factors, the best guide RNA pairs are ranked.

In the case of CIITA exon 3 and CD52 exon 1 gene editing, a stop codon was added to the beacon template on the forward atgRNA to prevent further translation. Overall, 24 spacer pairs targeting the three genes were selected and screened experimentally, and the top-performing pairs based on beacon placement efficiency were chosen for our cell therapy program.

For example, at the CD52 locus, we designed a total of 18 dual-atgRNA pairs to target Exon 1 and Exon 2. As shown in Figure 1A, these pairs included combinations of dual atgRNAs designed to maximize cleavage and beacon integration efficiency. Beacon integration efficiency was measured using droplet digital PCR (ddPCR) at the CD52 locus for each atgRNA pair. Among the 18 tested atgRNA pairs, the AA2905 atgRNA pair, targeting Exon 2, demonstrated the highest beacon integration efficiency at 45% (highlighted by an arrow in Figure 1B). This efficiency was notably higher than other pairs, making AA2905 the preferred candidate for subsequent validation and functional studies.

**Figure 1.**
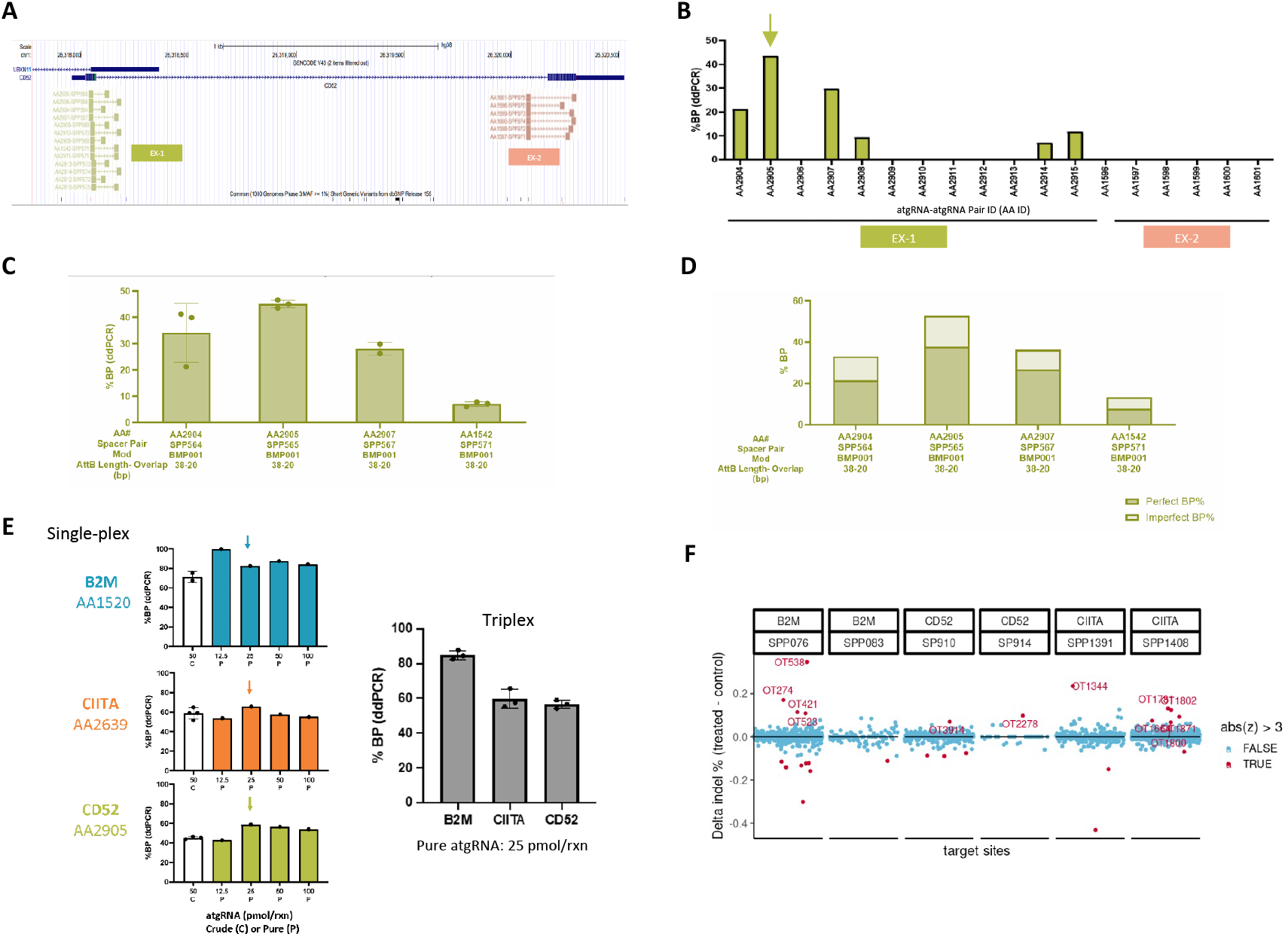
Beacon Placement in iPSCs. **A)** Representative CD52 atgRNA screen. Schematic representation of 12 dual atgRNA pairs targeted at Exon 1 and 6 atgRNA pairs targeted at Exon 2 for CD52 locus. **B)** Beacon integration by ddPCR at CD52 locus. AA2905 atgRNA pair yields highest ∼45% beacon placement efficiency (highlighted by arrow) among the 18 atgRNA pairs tested. Beacon integration by **C)** ddPCR and **D)** Amp-seq for top CD52 atgRNA confirms AA2905 yields the highest beacon placement efficiency. **E)** Singleplex and Triplex beacon placement in iPSCs. Left: Bar plots depicting the dual atgRNA titration with top selected atgRNA pairs at B2M, CIITA and CD52 loci. Purified atgRNAs yield highest beacon placement efficiency. Right: Triplex beacon placement in iPSCs with top atgRNAs (highlighted by arrows in left panel) yields beacon placement efficiencies of ∼90%, ∼60%, ∼60% obtained at B2M, CIITA, CD52 loci. **F)** Off-target evaluation. Tested *in silico CALITAS* predicted off targets for 5 atgRNA pairs at B2M, CIITA and CD52 for cellular validation. Detected possible off-target indels at 2/1,541 analyzed sites using rhAMP-seq.

To confirm the ddPCR results, we validated beacon integration for the top 4 atgRNA pairs using ddPCR and next generation sequencing (NGS). As illustrated in Figures 1C and 1D, ddPCR and NGS data confirmed that the AA2905 atgRNA pair yielded the highest beacon integration efficiency among all tested pairs. Specifically, AA2905 demonstrated a robust and reproducible integration profile at the CD52 locus, affirming its suitability for downstream applications in iPSC editing and providing a reliable marker for successful genomic modifications.

After identifying the AA2905 pair as optimal for CD52 targeting, we extended our analysis to evaluate beacon placement efficiencies for two additional target genes: B2M and CIITA. We performed beacon placement in both singleplex (individual dual atgRNAs for each target) and triplex (combined dual atgRNAs for three targets simultaneously at different atgRNA concentrations) configurations to assess multiplexing potential in iPSC editing (Figure 1E). In the singleplex configuration, each target demonstrated high beacon integration efficiency at 25 pmol/reaction of each gRNA, achieving approximately 80% for B2M, 60% for CIITA, and 66% for CD52 (left panel of Figure 1E). This high efficiency in singleplex conditions indicated robust integration capability for each gene individually.

For triplex beacon placement, we combined each of the atgRNAs at 25 pmol targeting B2M, CIITA, and CD52 to evaluate simultaneous editing efficiency. Triplex editing yielded comparable integration efficiencies to singleplex conditions, with each target achieving similar integration rates in the multiplex setup (right panel of Figure 1E). These results highlight the feasibility of performing multiplex gene editing with minimal compromise on beacon integration efficiency, potentially enabling multi-target editing in iPSCs for more complex genetic engineering applications.

Given the potential for off-target effects in CRISPR-based editing, we performed *in silico* analysis using the CALITAS tool to predict possible off-target sites for the selected gRNAs at B2M, CIITA, and CD52 loci. This analysis identified a set of potential off-target sites for each gRNA, which were further assessed experimentally using rhAMP-seq across 2,154 genomic regions (Figure 1F). The analysis revealed that off-target integration events were minimal, with low deviation from expected indel frequencies across all tested loci. Most off-target events exhibited indel counts well within acceptable limits, as shown in Figure 1F, where only a few sites showed more than three deviations from the median across all loci.

### I-PGI in iPSCs

After optimizing the beacon placement, we conducted a comprehensive screen of electroporation parameters to maximize I-PGI efficiency while maintaining cell viability in iPSCs. Various voltage and pulse width conditions were tested with an EF1a promoter-driven GFP plasmid as a tool cargo. Our results revealed that a 1500V, 20 ms single pulse yielded optimal results, achieving a balance of 53% cargo integration by ddPCR (indicating successful integration) and 40% cell viability (Figure 2A). This condition was consistently applied in subsequent experiments to ensure reproducibility and maximize I-PGI efficiency.

**Figure 2.**
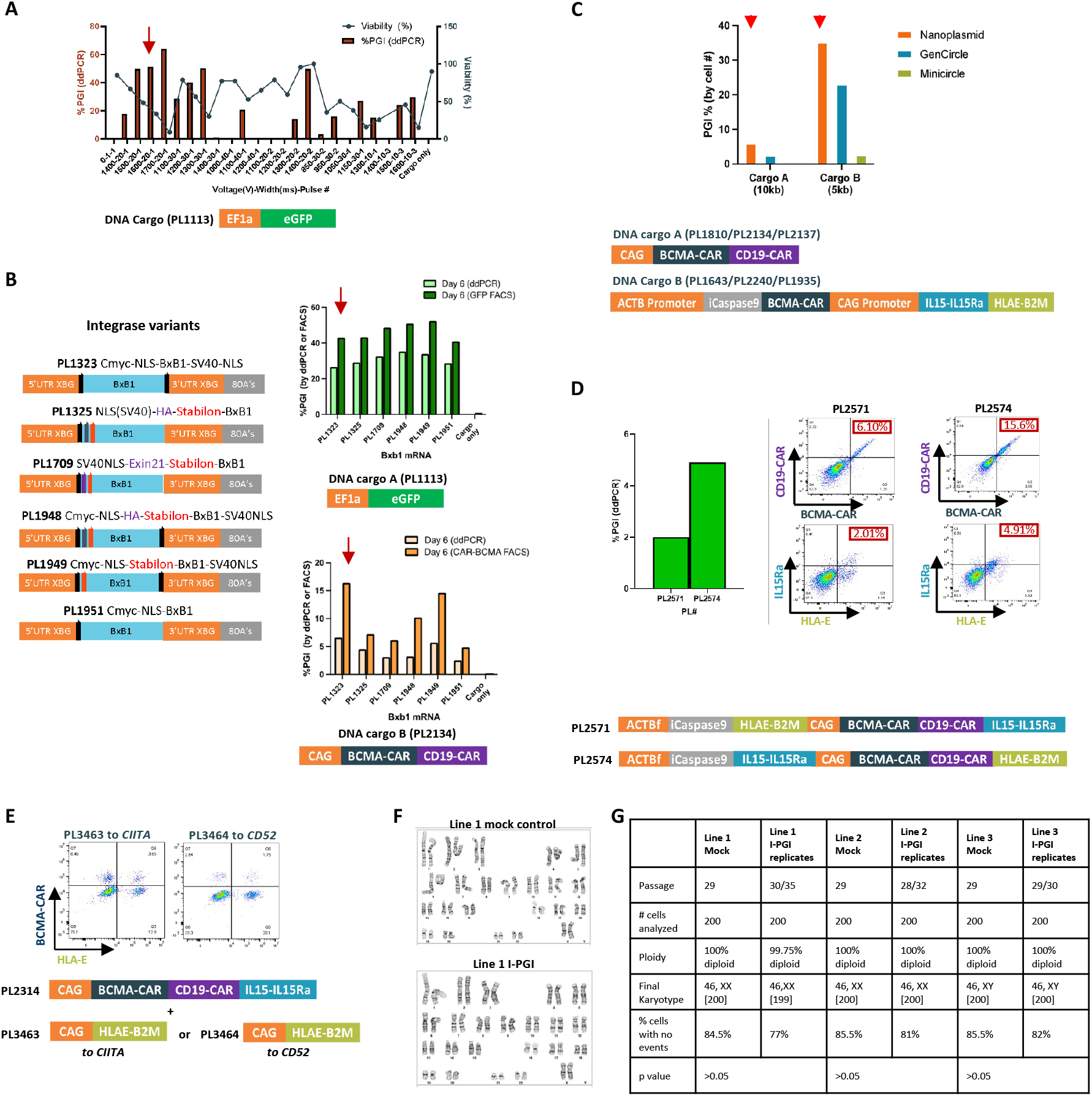
Integrase-mediated programmable genomic integration in iPSCs. **A)** Electroporation code optimization for PGI. PGI with 24 different electroporation programs identified 1500V-20 ms-1 pulse (highlighted by arrow) as the optimal electroporation program yielding ∼55% PGI and ∼40% viability 3 days after electroporation. EF1a promoter driven GFP plasmid used as the template cargo. **B)** Comparison of different Bxb1 integrases for PGI. Left: Schematic representation of different Bxb1 integrase versions tested for PGI in iPSCs. Elements evaluated in the variants include C-myc and SV40 NLSs, Stabilon and Exin21 stabilizing sequences, and human influenza hemagglutinin (HA) tag. **Right:** C-myc and SV40 NLS Bxb1 integrase yields highest PGI efficiency (highlighted by arrow: ∼8% by ddPCR, ∼16% CAR-BCMA by flow cytometry on Day 6) among the 6 tested Bxb1 integrase mRNAs. CAG promoter driven dual BCMA-CD19 CAR nanoplasmid used as the template cargo. **C)** Comparison of different cargo formats for PGI. Nanoplasmid shows highest cargo integration followed by gencircle with two different cargos: cargo A: Ef1a driven GFP; and cargo B: ACTB promoter driven iCaspase9, CAR-BCMA; and CAG promoter driven IL15, HLA-E. **D)** Integration of long DNA cargo. Successful insertion of cargo up to 16kb kb full-length cargo demonstrated by ddPCR which drives robust transgene expression as depicted by protein expression by flow cytometry. **E**) Successful multiplexed integration of two different cargoes at B2M and CD52 as demonstrated by protein expression by flow cytometry. **F)** Representative karyotypic analysis of iPSCs after integrase-mediated programmable gene insertion showed normal karyotypes indicating no major chromosomal abnormalities. **G)** Karyotyping data table summarizes the analysis of 200 cells from each condition, with n=2 lines from the I-PGI conditions, supporting no significant chromosomal abnormalities from I-PGI.

We next assessed multiple Bxb1 integrase variants to identify the configuration with the highest efficiency for I-PGI in iPSCs. These variants were designed with different combinations of nuclear localization signals (NLSs), including C-myc NLS and SV40 NLS, and stabilizing sequences, such as stabilon and Exin21^20,21^, all aimed at increasing the integration by Bxb1.

The I-PGI efficiency of each variant was measured by ddPCR and by quantifying GFP or CAR-positive cell populations using flow cytometry at Day 6 post-integration. Among the constructs tested, the variant Cmyc-NLS-BxB1-SV40-NLS (PL1323) consistently demonstrated the highest I-PGI efficiency across both cargos (Figure 2B), likely due to the enhanced nuclear localization and stability of the Bxb1 integrase, contributing to higher efficiency in integration.

We next evaluated the impact of different cargo formats on I-PGI efficiency. We compared the integration efficiency of nonviral circular templates, including Nanoplasmid, GenCircle, and minicircle vectors, each encoding the GFP gene (small cargo) and larger constructs (encoding antigen-specific targeting BCMA-CAR and CD19-CAR). Results indicated that the Nanoplasmid format consistently achieved the highest levels of integration across both small and large cargos, as reflected by the ddPCR integration and percentage of GFP-positive cells post-electroporation (Figure 2C). Nanoplasmid constructs also demonstrated superior performance with larger constructs, including BCMA-CAR and CD19-CAR. This finding suggests that Nanoplasmid-based vectors offer enhanced stability and integration efficiency, and were therefore selected for use in subsequent experiments involving complex genetic constructs.

To explore the capability of I-PGI for therapeutically relevant large sized gene insertion, we attempted integration of two different constructs of ∼13.5 kb size, encoding BCMA-CAR and CD19-CAR, IL-15Rα, HLA-E and inducible Caspase 9 (iCasp9) at the B2M locus in iPSCs. Post-electroporation, flow cytometry analysis confirmed co-expression of both CARs, with a distinct population of double-positive cells indicating successful integration. Specifically, the PL2571 construct yielded 6% BCMA-CAR and CD19-CAR double-positive cells, while PL2574 resulted in 15.6% double-positive cells (Figure 2D). These data demonstrate the feasibility of using I-PGI for large cargo insertion to engineer iPSC lines expressing multiple transgenes, a capability with potential applications in creating multifunctional therapeutic cell lines.

To demonstrate the feasibility of simultaneous multi-gene insertion using I-PGI, we introduced two constructs, one encoding BCMA-CAR, CD19-CAR, and IL-15Rα into B2M locus and the other encoding B2M-HLA-E, key component for immune evasion, into CIITA or CD52 locus.

Flow cytometry analysis confirmed the stable expression of HLA-E and BCMA-CAR, with successful detection of 3.65% and 1.75% HLA-E /BCMA-CAR double-positive cells in constructs PL3463 and PL3464, respectively (Figure 2E). These results confirm that I-PGI can achieve targeted integration for functional immune-related genes in iPSCs, paving the way for the generation of hypo-immunogenic cell lines for allogeneic applications.

To further assess whether I-PGI impacts genomic integrity, we conducted karyotype analysis on I-PGI-edited iPSCs from 3 independent lines. For each parental line, 2 technical replicates of I-PGI-engineered cells and 1 unedited control were analyzed. Metaphase spreads analysis revealed no statistically significant chromosomal abnormalities in I-PGI-edited iPSCs compared to unedited control lines, indicating that I-PGI did not compromise genomic stability (Figures 2F, 2G). This result is critical for validating the safety of I-PGI as a reliable and stable method for therapeutic genome editing. The absence of chromosomal abnormalities supports the potential of I-PGI for clinical applications, where maintenance of karyotypic integrity is critical.

### Single cell clone production and screening

After genomically engineering the iPSCs using I-PGI, we developed a highly efficient workflow to select single cell clones with desired on-target editing (Figure 3A). We first incorporated a staining and enrichment step with single cell deposition utilizing the Namocell Pala instrument. This instrument was chosen as it uses gentle and low-pressure deposition for cell sorting, which reduces stress on the iPSCs. By flow cytometric staining the I-PGI’d iPSC population for one of the proteins of interest, we enriched and deposited single cells expressing the stained protein from the I-PGI cargo into 96-well plates. We then allowed the cells to recover and clones to grow, and single-cell passaged the cells into two 96-well plates, one plate used for DNA extraction and genomic analysis, and the other plate cryopreserved for banking. A multiplexed amplicon sequencing, Reverse-transcription High-fidelity Amplification and Sequencing (rhAmpSeq), was used to identify clones with desired editing, including biallelic beacon placement at all 3 loci (B2M, CIITA and CD52), and mono– or bi-allelic cargo insertion into the B2M locus.

**Figure 3.**
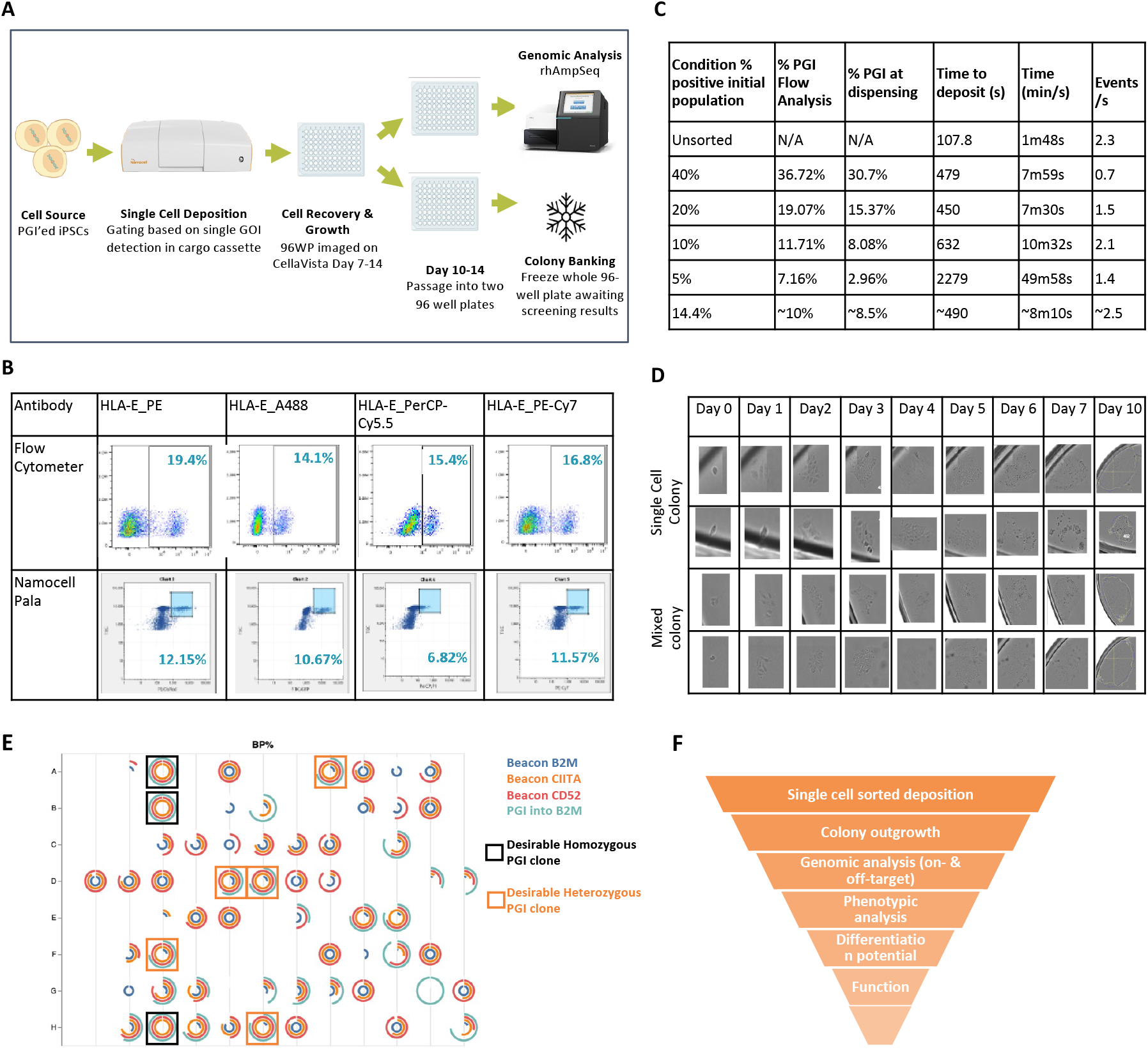
Single Cell Sorted Deposition and clone screening. **A)** Schematic representation of the workflow for single cell sorted deposition using the Namocell Pala Instrument for single cell sorted deposition, followed by clone banking and genomic analysis by rhAmpSeq. **B)** Representative plots from the fluorophore screening comparing the same stained samples run on both the Namocell Pala and the Cytoflex LX; PE shows the best separation on the Cytoflex and subsequently the Pala. **C)** The deposition time was monitored using different %PGI of the initial population while controlling events/second. The lower the %positive cells, the longer it take for the plate to deposit, impacting viability of the clones. **D)** Representative images of clonal assessment using the Cellavista, with two examples of a single cell colony and two of a mixed clonal colony. **E)** A rhAmpSeq plot of a representative 96 well plate analyzed to select genetically desirable clones; examples of homozygous PGI clones are boxed in black and examples of heterozygous PGI clones are boxed in orange. **F)** Summary of clone screening process. Post single cell outgrowth and on-target genomic analysis described, further assays, including phenotypic analysis, differentiation of iPSC into target cell type, functional assessment, and off-target and genome integrity evaluation, can be performed to identify the master cell clone for downstream application.

We first sought to optimize the sorted single cell deposition by evaluating different antibodies conjugated with fluorophores (Figure 3B). HLA-E was used as our protein of interest expressed by the I-PGI cargo. Among the same antibody clone conjugated to 4 different fluorophores detectable by the Pala, PE fluorophore resulted in the best separation between positive and negative populations and the closest percentage of positive cells compared to the traditional flow cytometry instrument, CytoFlex LX (Beckman Coulter). HLA-E_PE was thus chosen in the sorted cell deposition workflow.

We next assessed the feasibility of the sorted single cell deposition with different starting iPSC populations containing varying levels of I-PGI. The speed of deposition in the Pala was controlled to balance minimal doublet cell deposition into a single well, and to prevent prolonged dwelling time for optimal cell health. We started with an iPSC population with an initial I-PGI of 40%, and diluted it with unedited cells to 20%, 10%, and 5%, as well as a bulk edited population of 14.4% I-PGI, and a non-sorted deposition for control (Figure 3C). Different parameters were monitored during the workflow, including the %I-PGI detected by flow cytometry (CytoFlex LX) after dilution, the %I-PGI detected by Pala, the time of deposition for one plate (averaged from 2 plates), and the events/sec rate. We observed a significantly prolonged time of deposition when the starting I-PGI percentage decreased from 10% to 5%, which negatively impacts cell health and clonal outgrowth. Therefore, we established 5% starting I-PGI in iPSCs as a feasible threshold for successful sorted deposition.

Furthermore, we monitored the outgrowth and clonality from the 96-well plates using the CellaVista instrument from Synentec. This instrument is an automated ultrahigh-throughput imaging system that non-invasively measures brightfield and fluorescence images from the 96-well plates. Images were taken every day for a week and a final image at day 10, with the day 0 image taken 2-4 hours after deposition to allow the cells to settle and attach. Images of 2 single cell colonies and 2 mixed colonies are shown in Figure 3D. Such longitudinal imaging during the iPSC single cell outgrowth phase facilitated the selection of clones without mixed colony contamination.

After the outgrowth of single cell clones, rhAmpSeq, a high-sensitivity multiplex sequencing technology, was performed to evaluate beacon placement at the three loci of interest (B2M, CIITA, and CD52), and I-PGI at the B2M locus. An in-house bioinformatics tool was developed to analyze the data from each clone and plot it using 4 concentric circles as shown in Figure 3E as one example plate. From inner to outer circle, the orange circle represents BP at CIITA, red represents BP at CD52, blue is BP at B2M, and teal is I-PGI at B2M. The completeness of each circle represents the % alleles edited, which in a single clone would be 50% or 100%; any deviation could indicate either a mixed clone or a bias of the sequencing assay. Desirable clones with I-PGI in B2M, either homozygous or heterozygous with BP in the other allele, and biallelic BP in CIITA and CD52 loci are highlighted.

Following the identification of clones with desirable on-target editing, the corresponding wells from the banked plates can be thawed and expanded. Further analysis, including phenotypic characterization, differentiation of iPSC to target cell type, functional assays, off-target and genomic integrity assessment, can be performed to identify the master clone suited for the downstream use (Figure 3F).

## Discussion

In summary, our data demonstrate the ability to successfully screen for atgRNAs with high beacon integration efficiency in iPSCs, as we have outlined with the choice of AA2905 atgRNA pair for the CD52 locus. Additionally, BP efficiency is maintained in the triplex configuration compared to singleplex, as demonstrated by the triplex BP at B2M, CIITA, and CD52 loci, supporting the versatility of our approach and further showcasing I-PGI as a high-efficiency gene editing tool for multiple targets within iPSCs. The minimal off-target integration, as validated by rhAmpSeq, further underscores the specificity and reliability of our optimized atgRNA pairs.

Furthermore, we optimized parameters for efficient I-PGI in iPSCs, including electroporation conditions, integrase selection, and DNA cargo format, achieving a high level of targeted genomic integration with preserved cell viability and stability. By utilizing a Bxb1 integrase variant and nanoplasmid cargo, we obtained robust I-PGI efficiencies for both single and dual transgene insertions, as demonstrated by successful integration of GFP, BCMA-CAR, CD19-CAR, HLA-E-B2M constructs. The ability to target specific genomic loci while preserving karyotypic integrity establishes I-PGI as a powerful tool for complex genetic modifications in iPSCs.

Notably, I-PGI shares its core mechanistic foundation with PASTE and PASSIGE technologies, all of which utilize a CRISPR-directed nickase fused to a reverse transcriptase to install a recombinase or integrase recognition site (the “beacon”) followed by site-specific integration mediated by a large serine integrase such as Bxb1. Thus, these technologies represent conceptually equivalent strategies for double-strand-break-free genomic insertion, differing primarily in construct design and implementation.

We then established an efficient workflow for single cell clone production and screening for a master cell clone. Our streamlined process coupled single cell deposition with an enrichment step by staining and selecting for cells expressing I-PGI’d transgenes on the cell surface. We then tracked clonality with longitudinal imaging, and developed a high throughput screening process using rhAmpSeq to identify clones with intended KIs and KOs. The selected clones with desirable multiplexed genome engineering can be further interrogated for differentiation and functional assessment. This workflow provides the iPSC-derived cell therapy industry with a quick manufacturing of master cell clones from highly edited iPSCs while decreasing timelines and costs, two major constraints in the cell therapy production industry.

## Conclusions

We present here a highly efficient genome engineering process in iPSCs using I-PGI technology, followed by a rapid clone generation and selection process to identify clones with intended editing. Our novel approach significantly improves the speed and precision of CRISPR-based genome editing in iPSCs, paving the way for advanced cell engineering applications. Our findings also highlight I-PGI’s potential for advancing the development of next generation cell therapies with broad applications in regenerative medicine and immunotherapy.

## Supplemental Tables

Table 1. ddPCR and NGS primers

Table 2. List of antibodies

## Author Contributions

K.M. and R.H. conceptualized the study, performed the experiments, analyzed the data, and wrote the manuscript. M.Z., A.M.B., M.B., B.C.O., C.M., M.K., L.H., A.B. A.A., M.F., D.H., D.J.O., D.A., M.G., J.G., X.S., D.H., S.K., R.A., contributed to the wet experiments, J.W., D.N., D.S., W.W. performed the bioinformatic analysis. J.D.F supervised the study. C.L. conceived the study and wrote the manuscript.

## Acknowledgments

We would like to thank all members of Tome Biosciences for their support in the generation of this work.

